## Supplementary material for "A new pipeline SPICE identifies novel JUN-IKZF1 composite elements": Table 1

| STATs (Conditions) | Identified canonical GAS motif | GAS% | Tetramer likelihood | Optimal Spacing (bp) |
|---|---|---|---|---|
| STAT1 (MΦ, +IFN-γ) | 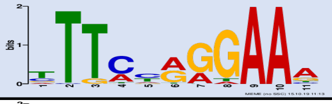 | 87.4% | **** | 10 |
| STAT2 (Th1, +IFN-γ) | 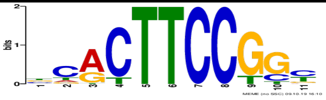 | 38.2% | NA | NA |
| STAT3 (CD8T, +IL-21) | 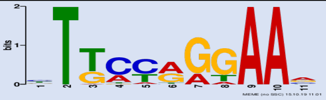 | 87.0% | *** | 11? |
| STAT4 (Th1) | 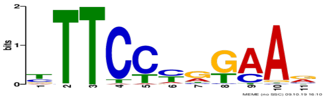 | 95.4% | **** | 11-12 |
| STAT5A (Total T, +IL-2) | 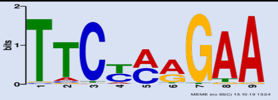 | 99.8% | **** | 6-7, 11-12 |
| STAT5B (Total T, +IL-2) | 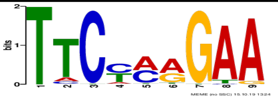 | 100% | **** | 6-7,11-12 |
